# A Population Model Reveals Surprising Role of Stochastic Cell Division in Epigenetic Memory Systems

**DOI:** 10.1101/2025.03.17.643620

**Authors:** Viviane Klingel, Dimitri Graf, Sara Weirich, Albert Jeltsch, Nicole E. Radde

## Abstract

Epigenetic memory systems can store transient environmental signals in bacteria in form of DNA methylation patterns. A synthetic zinc finger protein (ZnF4) binds to the DNA in a methylation-dependent manner and represses the expression of the DNA methyltransferase CcrM. The ON-state of these systems is characterized by high CcrM expression, high methylation levels, and low ZnF4 binding, but the mechanisms ensuring long-term ON-state stability remain unclear. Measurements showed a gradual shift of cell populations from ON to OFF starting after about four days of cultivation. We use a hybrid modeling approach integrating flow-cytometry data and bulk methylation measurements to test the hypothesis that stochastic cell division is a key factor in this transition. Interestingly, model parameters cluster into two groups with opposite effects of cell division rates on ON-state stability. Experiments under varying growth conditions show that faster cell division increases memory stability — an initially unexpected result. Model simulations provide a potential explanation for this observation and deepen our understanding about the mechanisms and timing of the ON/OFF switch in individual cells.

## 1 INTRODUCTION

Epigenetic information can be stably inherited during cell division, but can also be reverted, as no genetic changes occur (Berger et al. 2009). Both properties make epigenetic processes attractive targets for the design of synthetic sensor and memory systems (Maier et al. 2017, Khalil and Collins 2010). In recent years, different synthetic epigenetic systems that use DNA methylation as a key epigenetic signal have been designed and implemented in bacteria and other organisms (Fernández-Fernández et al. 2024, Maier et al. 2017, Park et al. 2019, Van et al. 2021). Specifically, we are here interested in synthetic epigenetic memory systems described in Maier et al. (2017) (Figure 1) and derived systems (Ullrich et al. 2020, Graf et al. 2023). Their key player is an engineered zinc finger protein (ZnF4), an artificial DNA binding protein that binds in a methylation-dependent manner to the DNA and thereby represses the expression of the DNA methyltransferase (CcrM). The system is initially in a stable OFF-state, which is characterized by low CcrM expression, low DNA methylation and tight, cooperative binding of ZnF4 to the DNA (Figure 1a). The memory plasmid is combined with a trigger plasmid allowing the induction of CcrM expression by addition of arabinose to the growth medium (Figure 1b), which is detected as an mCherry signal. If enough trigger CcrM is present, this can methylate the DNA in the CcrM promoter of the memory plasmid, which prevents ZnF4 from binding, and induces the ON-state of the system. In the ON-state, CcrM expression is high, and ZnF4 binding to the DNA is repressed by high methylation. The ON-state persists even if expression of the trigger CcrM is stopped by addition of glucose (Figure 1c), finally allowing the system to memorize transient input signals.

**FIGURE 1.**
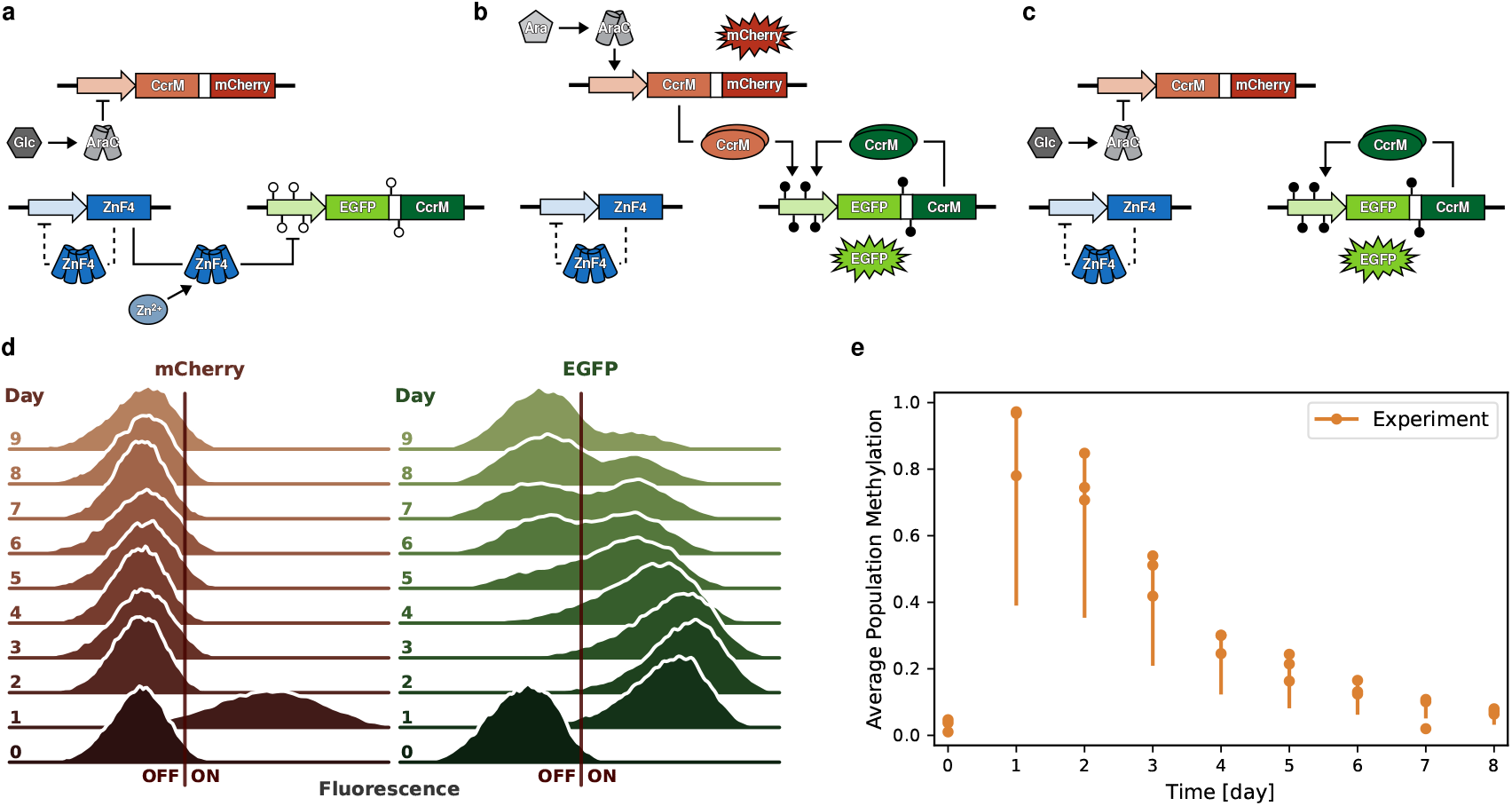
Biological Memory System. **a–c:** The schemes show three states of the system (modified from Maier et al. (2017)): **a**: OFF-State: In the OFF-state ZnF4 represses CcrM expression, therefore methylation levels are low. Expression of the trigger plasmid is repressed through AraC in the presence of glucose. The OFF-state can be stabilized or induced by increasing the Zn^2+^ concentration, which leads to stronger ZnF4 binding (Graf et al. 2023). **b:** Triggered ON-state: The ON-state can be triggered by switching from glucose to arabinose, which leads to the expression of CcrM and mCherry from the trigger plasmid. CcrM can then bind and methylate in the regions that are bound by ZnF4, which has a negative effect on the binding affinity of ZnF4, thus weakening its repressive effect. This turns on a positive feedback loop through the following expression of CcrM from the memory plasmid. EGFP and mCherry are used to detect the expression of CcrM from both plasmids. **c:** ON-state: After one day, the trigger is turned off by switching back to glucose. CcrM and EGFP from the memory system continue to be expressed, while enough CcrM and therefore methylation are present to inhibit ZnF4 binding. **d:** The experimental single-cell data from Graf et al. (2023) of flow-cytometry measurements of mCherry and EGFP fluorescence are shown as kernel density estimation on a logarithmic scale. The induction via the trigger plasmid can be observed as the increased fluorescence on day one for mCherry and slightly delayed for EGFP. mCherry returns to low fluorescence intensities after day one, while EGFP fluorescence remains high for a few days (memory effect) before slowly drifting towards the OFF-state. **e:** DNA methylation levels of the population are measured via restriction enzyme cleavage analysis. Due to the indirect nature of the measurement, the true methylation can be between 0.5–1x of the measured value (see Appendix Table S1). This range is indicated by the vertical bars. Data from Graf et al. (2023).

We have developed a data-based model to describe this epigenetic memory system which is able to recapitulate the bulk switching dynamics of CcrM and DNA methylation in bacterial cell cultures from the OFF to the ON-state upon different trigger inputs (Klingel et al. 2021). This model uses chemical reaction kinetics to describe the interrelated positive feedback loops between the molecular species in the system and operates in a bistable range with two stable steady states corresponding to the ON- and OFF-states of the system, respectively. This model provided retrospective insights into design steps of the memory system and was used to suggest an extended design for oscillations (Klingel et al. 2022).

However, this model is not suitable to provide insights into the long-term stability of the ON-state of the memory system, which was studied experimentally in detail in Graf et al. (2023) (Figure 1d). Upon induction of the ON-state via arabinose on day 1, the mCherry signal indicates a transient high CcrM level expressed from the trigger plasmid, which lasts for one day and was then turned off by switching from arabinose to glucose medium. The EGFP signal, which indicates CcrM expression from the memory plasmid, follows this induction with a slight delay and remains stable for three days.

After about four days of cultivation, the ON-state signal is gradually lost, which is caused by individual cells switching from the ONback into the OFF-state. Since switching times of individual cells in the population are highly heterogeneous, this gradual loss causes a slow drift of the population from high towards low CcrM levels lasting over several days. Approximately 3 days after removal of arabinose, the proportion of cells in the ON-state starts to decrease linearly with time, indicating that the probability of a cell switching into the OFF-state increases with the number of cell divisions (Graf et al. 2023). As expected, bulk DNA methylation levels of the population also gradually decrease during this drift (Figure 1e). Since both steady states are asymptotically stable in the deterministic bistable model, this model cannot account for this experimentally observed slow drift of the population towards the OFF-state. A simple simulation analysis, in which selected parameters of the model were varied, showed some heterogeneity at the population level, as for some parameter combinations the system was brought into a mono-stable region. In this region, only the OFF-state represents an asymptotically stable fixed point, but the system is still close enough to the saddle-node bifurcation point so that it only returns to the OFF-state after a residence time in the ON-state (Klingel et al. 2021). Since this residence time depends very sensitively on the parameters, a certain heterogeneity can be depicted with this approach, which, however, is not sufficient to explain the observed slow and reproducible drift towards the OFF-state.

A deeper understanding of the molecular mechanisms behind the experimentally observed drift behavior of bacterial cultures could be valuable to further optimize the design of more robust memory systems and applicable to related designed systems. To this end, we aim to develop a novel simulation and modeling approach to analyze this system, taking into account the decision making and switching mechanisms on a single-cell and on the population level. We theorize that the increasing probability of switching to the OFF-state might be due to an accumulation of stochastic effects over time. Our hypothesis is that stochasticity introduced by cell division is a major cause of cellular heterogeneity.

Cell cycle lengths vary over time and also differ between different cells. The main effect of cell division on our memory system is the halving of the DNA methylation, because during the cell cycle, new DNA strands are synthesized, which are initially not methylated. If, therefore, the concentration of CcrM is too low or if divisions happen too quickly, not enough new DNA methylation can be added, which is expected to weaken the stability of the ON-state. Further, proteins bound to the DNA, in our model ZnF4 and CcrM, are, at least temporarily, removed from the DNA during DNA replication. In the case of ZnF4, this can weaken the OFF-state because unmethylated ZnF4 binding sites, which previously were protected by ZnF4 binding, may become accessible to CcrM, leading to their potential methylation.

Here, we introduce a hybrid model which describes the molecular interactions between two cell divisions by reaction rate equations for a population of individual cells. Cell divisions are described by stochastic events which affect the molecular state of a cell. Our modeling takes into account that cell cycle lengths of mother and daughter cells are correlated. We use single-cell flowcytometry data of CcrM expression and DNA methylation measurements on the population level for model calibration. To this end, we formulate an inverse problem, which takes this multi-modal data into account and uses the Kolmogorov metric to compare cumulative distribution functions between real and simulated data. This metric is locatiON- and shape sensitive and is thus suitable to mimic a complex bimodal transient population behavior, as observed in our flow-cytometry experiments. The model is validated by using unseen experiments in which the memory system was induced into the ON-state and reset to the OFF-state several times in repeated cycles. We analyze the parameters and predictions of the validated model, which results in two competing scenarios about the role of cell cycle length on the stability of the ON-state. These scenarios are translated into the design of novel experiments which allow for selection between these scenarios. Surprisingly, these experiments clearly show that faster division tends to increase the stability of the ON-state. As our model predicts intra-cellular states as well as the population behavior, we can investigate the molecular mechanisms behind the experimentally observed properties of the OFF-switch and therefore gain valuable insights into the stability of the memory system.

## 2 RESULTS

### 2.1 Modeling Workflow

As the model design and calibration consist of many steps, we visualized our workflow in Figure 2. In the following we describe these steps in more detail and reference to the respective sections in this work. **1:** The model is designed based on knowledge about the underlying system and hypotheses or questions with regards to the experimental data it should represent (Section 2.2). **2:** An objective function is constructed to compare experimental data to simulation results. In our case, we used a distance measure for probability densities to compare the single-cell fluorescence data to the corresponding distributions of model states. Further we used penalties and constraints for indirect measurements like the DNA methylation data as well as qualitative information, like the known influence of the DNA methylation on ZnF4 binding (Section 2.3). **3:** The objective function is then used to optimize the unknown kinetic parameters in the model. This requires the choice of an appropriate optimizer and hyperparameters to account for the fact that the objective function varies slightly with each simulation even for fixed parameter sets due to the stochastic components of our hybrid model (Section 4.3). The observed behavior of the optimization as well as its success lead to revisions in the population model, for example to adjust the population size or sampled parameters (**I**). **4:** Parameters and simulation results obtained from an optimization have to be evaluated. The simulated states are compared to the expected biological behavior, supported by experiments, and to literature. The parameters are analyzed in terms of their identifiability as well as the robustness of the optimization (Section 2.4). Specifically, this has led to revisions of the objective function (**II**), for example by choosing another distance measure with different properties as well as the model itself (**III**), by for example considering different interactions or reactions, or reevaluation of previous model assumptions. This cycle of revisions was repeated many times until a satisfactory reproduction of the experimental data was achieved and we consider our model to be trained (**5**). We then challenged our model by simulating previously unseen experimental data (**7**), leading to a predictive model (**8**, Section 2.5). This then enabled us to generate and test different hypotheses of the underlying biological mechanisms and assumptions (**9**, Sections 2.6–2.7). The knowledge gained from confirming or disproving these hypotheses then led to further experiments and iterations of this modeling cycle (**11**, Section 2.7) which improved our understanding of invisible underlying processes.

**FIGURE 2.**
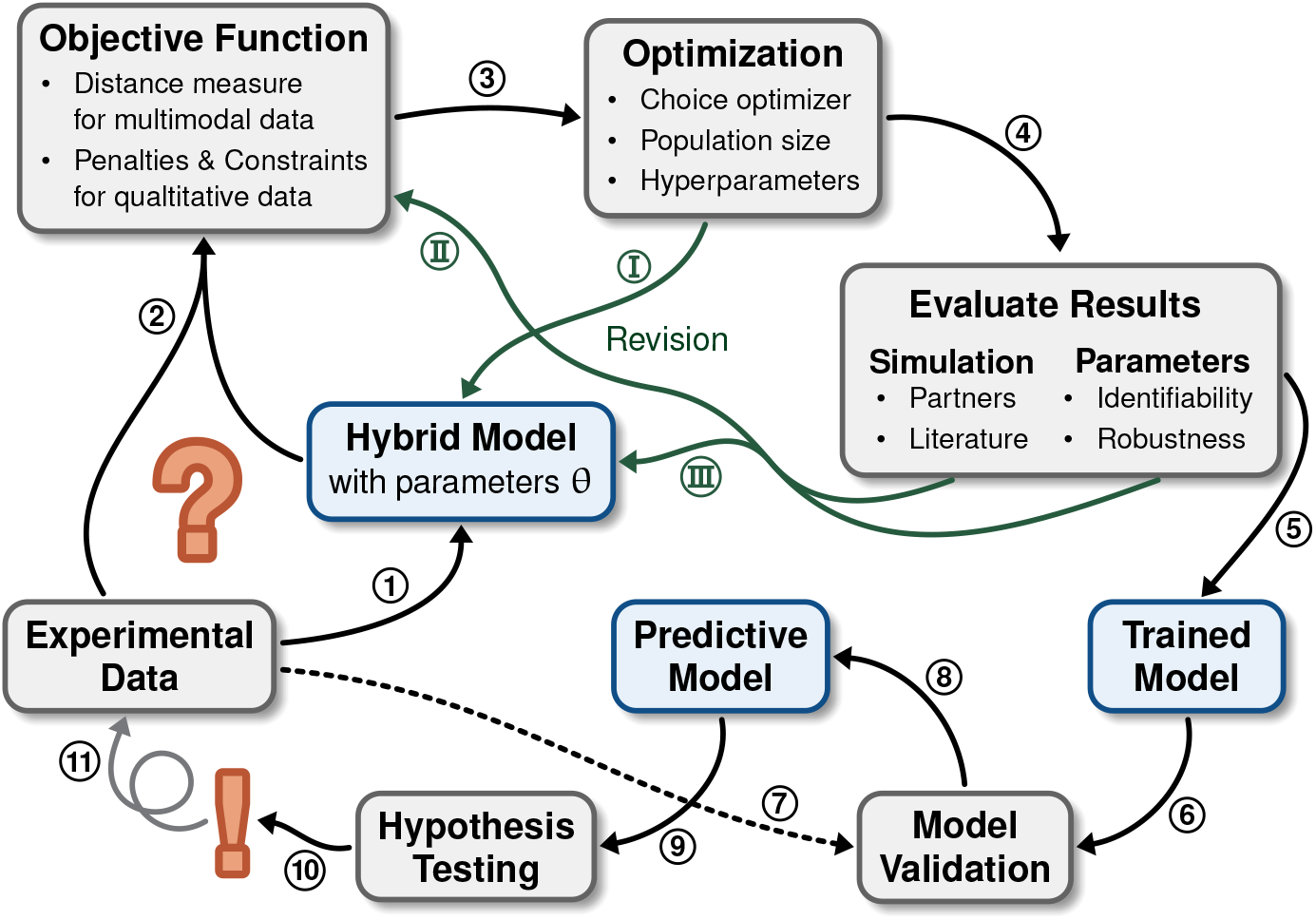
Workflow of model building and optimization. This workflow aims to illustrate some of the steps in constructing our hybrid model and its training to reproduce the experimental data, as detailed in Subsection 2.1.

### 2.2 Model Concept / Hybrid Simulation Scheme

Our hybrid population modeling approach has multiple layers, which are illustrated in Figure 3. With the aim of reproducing experimental dynamics of ON/OFF-state cell distributions and predicting single-cell trajectories, our model contains a population of individual cells. Each cell is simulated independently and in parallel. A possible approach to describe stochasticity and thus heterogeneity between different cells is a purely stochastic simulation. Our system, however, has large protein numbers (10^4^–10^5^) per cell, making a stochastic model inefficient and not very useful. Further, we had to solve an inverse problem, which is already quite complicated due to fitting a distribution rather than few population measurements. We have therefore chosen a hybrid simulation approach, consisting of deterministic biochemical processes at the individual cell level, which are interrupted by stochastic events caused by cell division. Our approach can be written as a stochastic ODE system for each cell,

**FIGURE 3.**
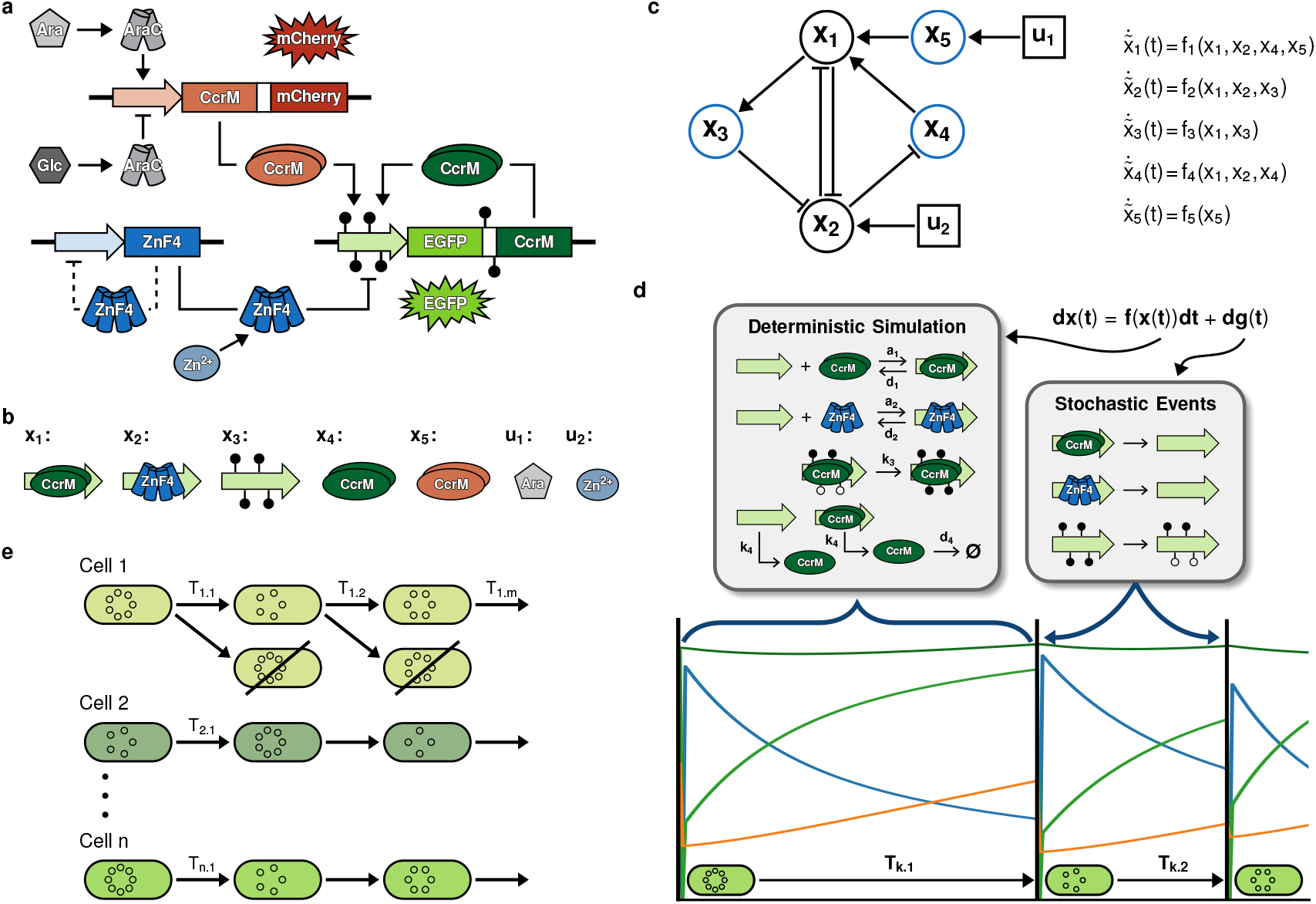
Illustration of Critical Features of the Population Model. **a:** The model scheme contains all possible interactions and states as described in Figure 1. Taken from Maier et al. (2017) in modified form. **b:** Translation of the different molecular species into mathematical states, including the two inputs used to switch the system ON or OFF. **c:** The interaction graph (left) of the deterministic part of the mathematical model (right) reveals the dependencies in the system including multiple feedback loops. Blue states indicate the states with experimental data. **d:** The hybrid simulation of single-cells is divided into two parts: I) A deterministic ODE describing protein binding, production and degradation, and DNA methylation. These correspond to the interactions shown in c. II) Stochastic events representing a simplified DNA replication and cell division, where proteins bound to DNA are removed and DNA methylation is halved. The time (*T*_*div*_) between two events is sampled from a random distribution, but allows for correlation within a cell to represent inheritance effects. Additionally, the number of plasmids at each division is also modeled as a random variable. **e:** The population model contains n individual cells, simulated separately and with different samples for cell cycle durations and plasmid copy numbers. Only one daughter cell after a cell division event is considered, such that the number of cells in the model remains constant. The resulting distributions of the model states can then be compared with the measured single-cell data.

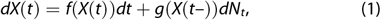

sometimes also denoted as a piecewise deterministic Markov process (Davis 1984). The term *f*(*X*(*t*))*dt* describes the flow of the system between events, i. e. the biochemical interactions of the memory system that constantly take place (Figure 3a): DNA binding and dissociation of CcrM and ZnF4, methylation of a methylation site, production and degradation of CcrM. The stochastic components of the model are captured by the term *g*(*X*(*t*–))*dN*_*t*_. While sharing the kinetic parameters that are optimized, heterogeneity between cells is achieved by randomly sampling plasmid copy numbers and cell division times.

#### Deterministic Model

The ODE model part is based on our previous work (Klingel et al. 2021) but further developed. Some of the changes include adaptations to account for the hybrid scheme as well as an altered description of DNA binding by CcrM, leading to a quadratic dependence of the binding rate on CcrM concentration, and methylation kinetics. Model states and inputs, as well as the model structure are visualized in Figures 3b and 3c, respectively. The ODE part of Equation (1) reads

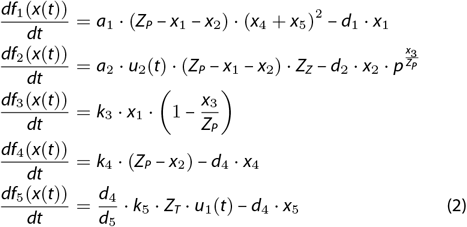

The states are listed in Table 1. The state *x*_1_ describes CcrM bound to its binding site on the DNA where it can methylate its target sequence. An unoccupied site is required for CcrM to bind at the rate *a*_1_. This is reflected by the term (*Z*_*P*_ – *x*_1_ – *x*_2_), where *Z*_*P*_ is the total number of plasmids in the cell, each with one binding site of interest. CcrM binds as a dimer (Horton et al. 2019), which may be formed by CcrM produced from either the memory plasmids (*x*_4_) or the trigger plasmids (*x*_5_). Bound CcrM dissociates at the rate *d*_1_. The state *x*_2_ corresponds to ZnF4 molecules that are bound to a binding site. The DNA binding of ZnF4 is sensitive to the methylation state of its binding site, with stronger binding when no methylation is present. We have modeled this as a methylation-independent association of ZnF4 to DNA and methylation-sensitive dissociation, where the influence of methylation is captured by the term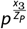. ZnF4 binding can be strengthened by an increased Zn^2+^ concentration, which is taken into account with the model input *u*_2_(*t*),

**TABLE 1.** States *x*_*i*_ of system (2).

|  |  |
| --- | --- |
| $x_1$ | CcrM bound to the DNA |
| $x_2$ | ZnF4 bound to the DNA |
| $x_3$ | Methylation state of a ZnF4 binding site |
| $x_4$ | CcrM from the memory plasmid |
| $x_5$ | CcrM from the trigger plasmid |

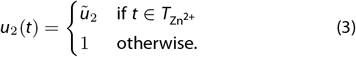

During the time interval with increased Zn^2+^ concentration,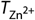, the ZnF4-DNA binding constant is increased by the factor*ũ*_2_. *Z*_*Z*_ corresponds to the concentration of free ZnF4, which depends on the plasmid number via *Z*_*Z*_ = 50 *· Z*_*P*_ (Graf et al. 2023). Between events this concentration is assumed to be approximately constant due to ZnF4 auto-regulation.

The state *x*_3_ describes the methylation state of a CcrM binding site. CcrM bound to the DNA (*x*_1_) can methylate a free methylation site with the rate *k*_3_. Following the biological setting of DNA-adenineN6 methylation in bacteria (Wion and Casadesús 2006), there is no active demethylation in this model. Hence the only mechanism for decreasing methylation is through DNA replication, where newly synthesized DNA strands are unmethylated.

The states *x*_4_ and *x*_5_ correspond to the concentration of CcrM derived from the memory or the trigger plasmids, respectively. If the promoter binding site is not occupied by ZnF4, CcrM is produced by the memory plasmids at the rate *k*_4_. ZnF4 binding blocks this transcription. CcrM from either source is degraded with the rate *d*_4_. CcrM from the trigger plasmids is produced during induction with arabinose, which is represented by the input *u*_1_(*t*),

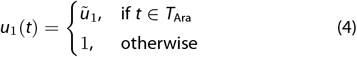

The induction strength is defined by the parameter *ũ*_1_, *T*_Ara_ denotes the duration of the arabinose induction. The parameter *Z*_*T*_ captures the heterogeneity in both trigger plasmid numbers and the strength of transcription induction or repression. As *f*_5_(*t*) only depends on variable *x*_5_ itself, we have chosen to optimize its parameters (*k*_5_ and *ũ*_1_) prior to the optimization of the full system to reduce the problem complexity and therefore save computation time. The time-scale of the subsystem used for this optimization step is coupled to the main system via the shared CcrM degradation rate *d*_4_. We therefore defined the subsystem

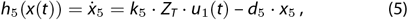

where the degradation rate, *d*_5_, is fixed during the optimization. In the full system, *h*_5_(*t*) is then normalized by the ratio 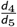, resulting in the expression for *f*_5_(*t*) in Equation (1).

#### Stochastic Cell Division Events

The deterministic simulation is interrupted by stochastic cell division events, where *g*(*X*(*t*–)) is a function describing the change due to the event and *N*_*t*_ are the number of events up to time *t*. The time between each event, *T*_*div*_, is randomly distributed and follows an exponentially-modified normal distribution. The events model the effect of DNA replication and cell division on the memory system (Equation (6)): With each replication, CcrM and ZnF4 bound to DNA are removed and the methylation level of the DNA is halved, as each newly synthesized strand is unmethylated. Furthermore, the number of plasmids changes at each division, depending on how many new plasmids are produced and how these are distributed among the daughter cells. We have expressed the stochasticity of plasmid numbers by a random variable (*Z*_*P*_) that follows a beta distribution, where the mean value corresponds to the experimentally determined average copy number of 160 memory plasmids per cell (Graf et al. 2023). The state change due to a stochastic event is given by

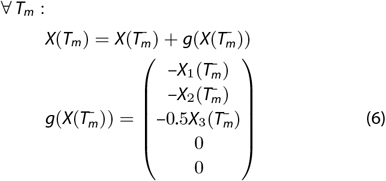

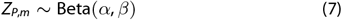

where *T*_*m*_ are the time points for a division and *X*(*T*^−^) is the last model state from the deterministic simulation before the division. The simulation scheme for an individual cell is illustrated in Figure 3d.

#### Model Outputs

To compare simulated values across cells with different stochastic parameters as well as to experimental data, we defined the following model outputs:

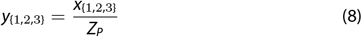

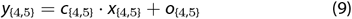

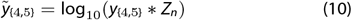

The states *x*_{1,2,3}_ are normalized to the number of plasmids *Z*_*P*_, resulting in ratios of bound CcrM and ZnF as well as the percentage of methylated sites. CcrM concentration is measured by its coproduced fluorescent reporters, mCherry for the trigger plasmid and EGFP for the memory plasmid. It is assumed that EGFP and mCherry are produced and degraded with the same rates as the corresponding CcrM, therefore they are considered to be equal to *x*_4_ and *x*_5_, respectively. The model outputs *y*_{4,5}_ correspond to the measured fluorescence intensities for EGFP and mCherry. *c*_{4,5}_ and *o*_{4,5}_ represent the parameters of a linear regression model that maps fluorescence intensities and the measured protein abundances for cells in the OFF-state (day 0) and in the ON-state (day 1). As these parameters are experiment-specific, they are calculated separately for each experiment (see Appendix Section 1, Figure S1). The states 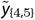 are used to compare the measured and simulated densities of fluorescence intensities on a logarithmic scale and include the normally distributed noise term *Z*_*n*_ (see Appendix Section 2).

#### Cell Population

The final simulated cell population consists of many individual cells, each with their own sampled values for the inter-division times and stochastic constants (Figure 3e). The number of simulated cells had to be chosen with some care. On the one hand, simulation times increase linearly with this number, which is especially critical for solving the inverse problem of parameter estimation, which requires repeated forward simulations of the population. On the other hand, since each cell is different, a sufficiently large sample is needed so that the population of simulated cells can represent the experimental data. In preliminary studies, we selected the number of cells so that an increase would not significantly change the outcome of the simulation. In our case, this condition was satisfied with *n* = 480 cells, which is a multiple of the number of parallel cores of a node at the high-performance computing facility we were using for running the parameter optimization. The number of simulated cells in the population remains constant over the simulation time, as we only follow the trajectory of one daughter cell (Figure 3e).

### 2.3 Objective Function and Optimization Results

Our objective function

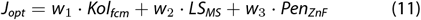

consists of three terms, which take different data types and system knowledge into account. The first term (*Kol*_*fcm*_) includes single-cell fluorescence intensities from flow-cytometry measurements. The second term (*LS*_*MS*_) accounts for bulk DNA methylation measurements, and the third term (*Pen*_*ZnF*_) accounts for the sensitivity of ZnF4 DNA binding to methylation levels. These terms are described in more detail in the following text. The weights *w*_1_, *w*_2_ and *w*_3_ are introduced to ensure appropriate relative scaling of each term.

#### Probability Metric for Single-Cell Data

For model calibration, we used experimental data taken from Graf et al. (2023). They provide flow-cytometry distributions of single-cells in ON- or OFF-states measured for ten days after triggering the system with arabinose and removal of the trigger, with three biological replicates each (Figure 1). We used a logarithmic scale for the fluorescence intensities for model calibration and visualization as these cover multiple orders of magnitude. Our aim in calibrating the model was to make the simulated densities as similar as possible to the experimental densities, with a particular attention to their shape. The shape is of relevance because of the multi-modal nature of the data. It represents the two subgroups of cells, namely those that are still in the ON-state and those that are in the OFF-state, with cells that are currently drifting into the OFF-state lying in between. In contrast, a collective OFF-drift of the entire population would appear as a distribution with a relatively constant shape moving towards lower fluorescence intensities.

Therefore, we paid special attention to choosing a probability metric for comparing experimental and simulated data that is sensitive to these shape differences. In our experience, the commonly used Kullback-Leibler divergence or Wasserstein metrics are inappropriate for this problem. Both sum or integrate over the entire support of the distribution and are thus less sensitive to rare but large differences. On the other hand, the Kolmogorov metric

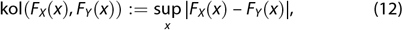

which is most commonly known as the test statistic used in the Kolmogorov-Smirnov test, considers only the maximum deviation between two distributions. Further, it is easy to compute because it only requires the absolute difference between the two empirical cumulative densities, without having to estimate the probability densities. It is also not very sensitive to a difference in sample sizes. The absolute difference over the entire data range is visualized for all three replicates of one day in Figure 4a. The maximum difference is marked by a star.

**FIGURE 4.**
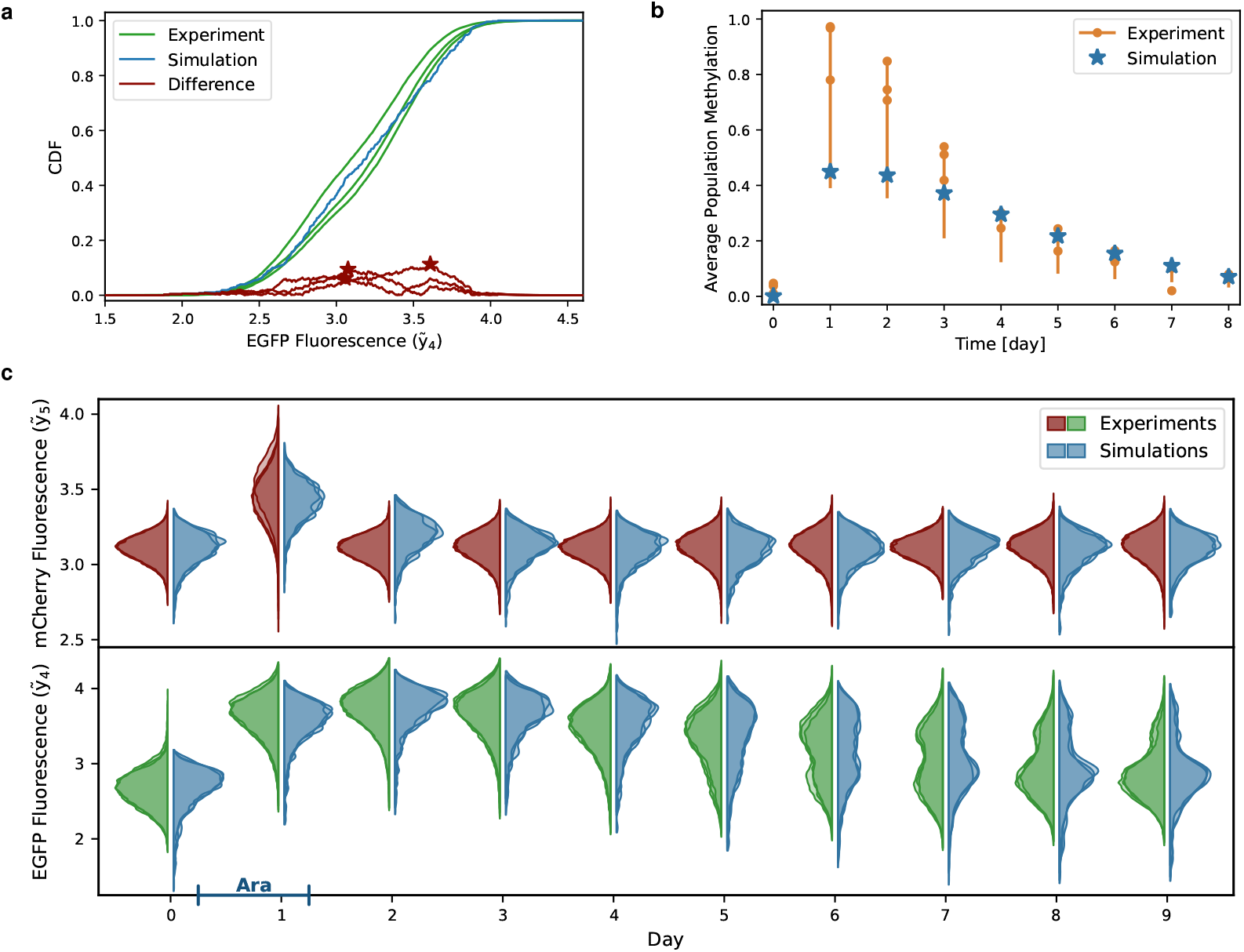
Optimization and model fit. Experimental data (Graf et al. 2023) and simulations from one representative parameter set. **a:** The main part of the objective function is a distance measure of the cumulative densities (CDF) between data and simulation of mCherry and EGFP fluorescence. The distance measure used here is the Kolmogorov metric, which is shape and location sensitive. The red stars indicate this metric as the largest difference between the CDFs of each of the data replicates and the simulation, which is used as a measure of distance in the Kolmogorov metric. CDF data are shown for one of the ten measured days. **b:** The indirect methylation measurements are included in the objective function with an adapted least-squares term. The measurements only give a range for the true methylation between 0.5–1x of the measured value. Therefore, any value within this range is not penalized in the objective function, every value outside receives the squared distance to the closest end of the range as a penalty. **c:** Violin plots of the measured and simulated densities of mCherry and EGFP fluorescence over ten days showing a very good fit of the model to the data. The probability densities are estimated using kernel density estimation. The data densities represent the three independent replicates, while the simulated densities were obtained from three different sets of randomly sampled values, one of which was used for the parameter estimation. The blue bar on the x-axis indicates the duration of arabinose induction, which activates the trigger plasmid leading to the mCherry signal and pushes the system into the ON-state. The fluorescence intensities in a and c are shown in a logarithmic scale.

The Kolmogorov metric was calculated for each replicate, day, and state, the sum of which gives us a measure of the total difference between simulated and experimental densities (*Kol*_*fcm*_).

#### Adapted Least Squares for Bulk Methylation Data

The bulk methylation state of the CcrM target region was measured using a restriction/protection-based DNA methylation detection system as described by Graf et al. (2023). The assay covers two restriction sites and the DNA is protected from cleavage if at least one strand of each site is methylated. To mimic the state after DNA replication, we assume that the two sites on a strand are either both methylated or both unmethylated. Therefore, if a site is protected from restriction, it can either be fully methylated or have one strand methylated and the other unmethylated. This results in a mapping of the true methylation state to the measured state, where both fully and half methylated sites are experimentally measured as methylated, while only fully unmethylated sites are assigned as such. So every population state between ‘all plasmids are fully methylated’ to ‘all plasmids are half methylated’, which corresponds to a true population methylation state between 50–100%, maps to a 100% methylation in the experimental restriction protection bulk measurement. To correct for this overestimation and partial uncertainty of the true methylation, we used a methylation range in the fitting process, assuming that the true methylation level lies between the measured value and half of this value. This range, represented by a vertical bar, and the original measurements are shown in Figure 4b. In our objective function this is implemented in form of an adapted least squares: Simulated methylation levels that lie within this range are not penalized. Values outside this range are penalized by a penalty equal to the squared distance to the nearest edge of the range. This is described by the methylation-dependent part of the objective function (*LS*_*MS*_).

#### Penalties for Qualitative Data

In the course of model building, optimization, and model analysis, we decided to add another penalty as part of the objective function that takes into account qualitative information about the memory system gathered during its design. A key design element of the system is ZnF4, which binds to the DNA in a methylation-sensitive manner, as described and quantified in Maier et al. (2017). Since we have no data directly characterizing the interplay between CcrM binding, methylation, and ZnF4 binding to the DNA, we first found several parameter combinations for which ZnF4 binding was insensitive to DNA methylation. Thus, we designed a penalty term *Pen*_*ZnF*_ that regularizes towards solutions that mimic the known sensitivity of ZnF4 DNA binding to the DNA methylation level, see also Section 4.3.

#### Optimization Results

We used a global optimization approach to search for optimal parameters and found several different parameter combinations with similar objective function values (Table 2, Appendix Table S2). Some of the parameters such as CcrM production (*k*_4_) and degradation (*d*_4_) rates as well as the methylation rate (*k*_3_) are relatively similar in all solutions and can thus be well determined. This is probably due to the fact that the EGFP fluorescence and the methylation data are very informative about these parameters. Other parameters such as ZnF4 and CcrM binding dynamics are less well identifiable, which is not surprising, since we do not have direct measurements of the molecular binding events at the DNA level. Interestingly, the ZnF4 association (*a*_2_) and dissociation (*d*_2_) rates clustered in two groups, with much less variation within the groups. We discuss these two groups in detail in Section 2.6. Overall, the best fitting parameter combinations behaved visually very similar in the model simulations (See Appendix Figure S2).

**TABLE 2.**
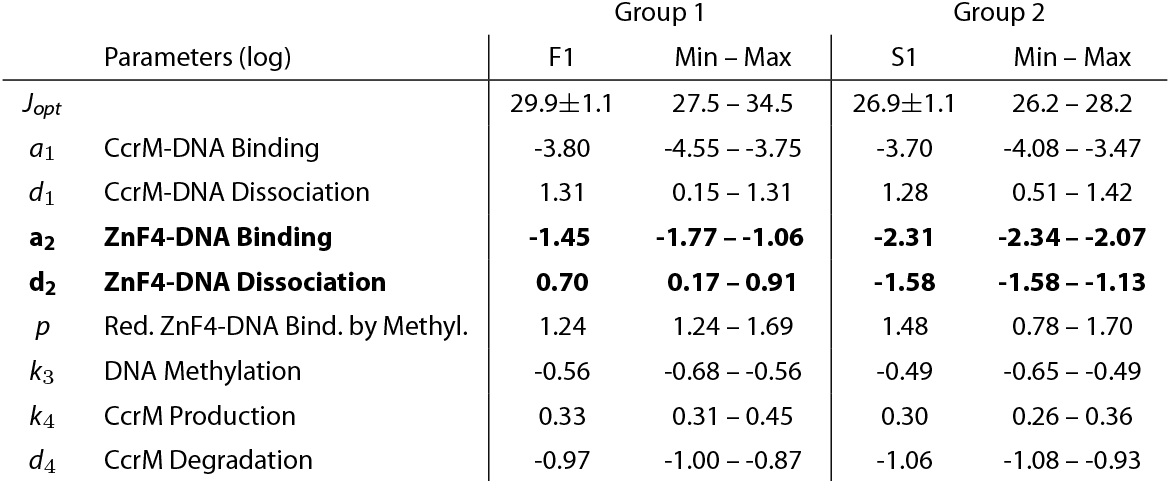
Parameter values. One exemplary set of parameter values for the observed two groups are shown together with their respective objective function value. The objective function value is the average ±the standard deviation of 50 individual simulations. Additionally, the minimal and maximal values from five different optimizations are given (See Appendix Table S2 for all parameter values). The values for the parameters displayed here are the logarithm of the parameters as they are used in the model.

### 2.4 Hybrid Model Captures Slow OFF-Drift

Our model was able to accurately reproduce the observed experimental data from Graf et al. (2023), as illustrated for one exemplary parameter combination in Figure 4. The empirical cumulative densities of an exemplary day show nicely how the simulated distribution matches the experimental ones (Figure 4a). The simulated average population DNA methylation levels also fall within the range provided by the measurements (Figure 4b). Both the time courses of simulated and measured methylation levels follow a slightly sigmoidal curve after day one, but with lower maximal values for the simulation. At high measured DNA methylation levels, the simulation is close to the lower end of the range, while at lower measured levels, the simulation is more towards the higher end. This could indicate that the actual methylation is in fact lower than previously assumed, while still maintaining the ON-state for many days.

The simulated densities of EGFP and mCherry fluorescence, which correspond to the CcrM levels expressed from the memory and the trigger plasmids, respectively, reflect the experimental measurements over the entire ten days well (Figure 4c). Here, we show three representative distributions of both the experiments and the simulations for each day. The experimental densities are the three independent biological replicates, while the simulated densities were produced by three different samples of distributed parameters and noise, which is sampled with every simulation. One of the experimental data sets was used during model calibration, the other two were used only for visualization. There are small differences between the three simulated densities, but they all show the same behavior and in many cases their variability is similar to that of the biological replicates. To analyze this observation quantitatively, we simulated each set 50 times and compared the resulting objective function values. We found only small to no differences between the three sets, and the distribution of objective function values roughly follows a normal distribution (Objective function values and histograms are shown in Appendix Tables S3, S4, S5 and Figure S3). We therefore conclude that the number of simulated cells is sufficiently large to avoid over-fitting to a specific set of distributed constants.

Importantly, while there are small discrepancies between model and experiment, especially in the tail behavior, the model overall captures well the slow OFF-drift of the population, and especially the bimodal nature of the EGFP fluorescence data. This is a significant improvement over our previous, deterministic model, which was unable to reproduce this slow drift of the entire population. Simulations in which we set either inter-division times or plasmid numbers to a constant value for each cell confirmed that stochastic cell divisions are mainly responsible for the slow OFF-drift, while varying plasmid numbers increase heterogeneity (Appendix Figures S6 and S7). Simulations in which both random variables were set to constant values are very similar to simulations of our previous work, where we had used a purely deterministic model with varying cell division rates (Klingel et al. 2021). Taken together, our model is able to accurately capture experimentally observed methylation levels and CcrM population dynamics, confirming our hypothesis that a stochastic process is required to fully describe these data.

### 2.5 Model Successfully Predicts Unseen Data

To validate our model, we used it to simulate previously unseen data, also from Graf et al. (2023). This includes one dataset that is a repeat of the experiments used for model calibration, and four experiments that employed a new mechanism of triggering a fast OFF-switching of the population (Figure 5). One of these experiments, in which the population is switched ON, OFF and ON again, is shown here (ON-OFF-ON), while the other two experiments (ONOFF and ON-OFF-ON-OFF) can be found in Appendix Figure S5. The measurement specific scaling factors for calculating protein amounts from fluorescence intensities were determined separately for these experiments (see Appendix Section 1).

**FIGURE 5.**
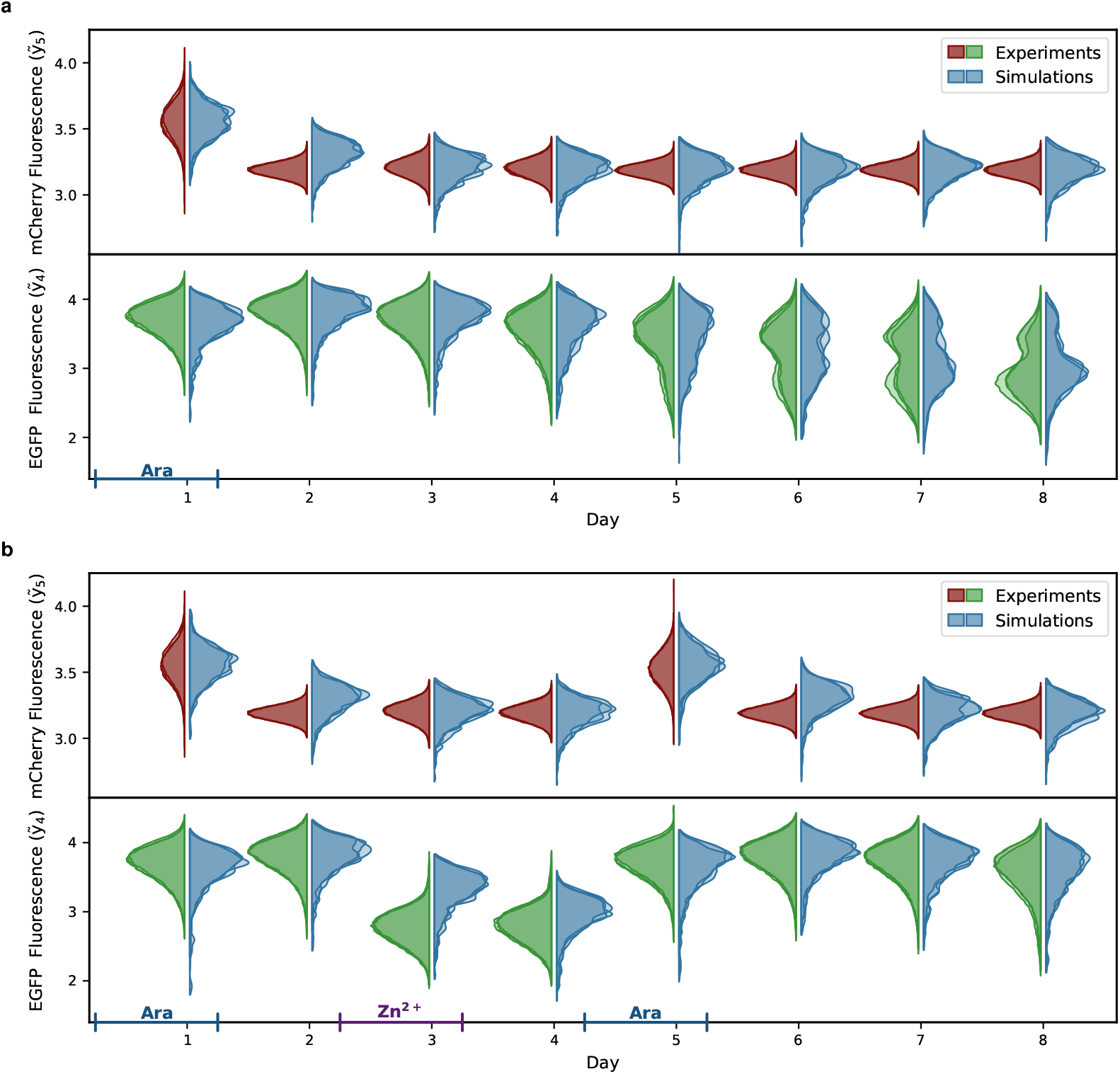
Model validation. Validation of the model with experimental data (Graf et al. 2023) not used in the optimization. Experiment and measurement specific parameters were adapted, specifically the scaling factors for calculating protein amounts from fluorescence intensities (Appendix Section 1). As in Figure 4, Violin plots of the measured and simulated densities of mCherry and EGFP over eight days are shown. The probability densities are estimated using kernel density estimation. The data densities represent the three independent replicates, while the simulated densities were obtained from three different sets of randomly sampled values, one of which was used for the parameter estimation. The blue and purple bars on the x-axis indicate the duration of arabinose induction or increased Zn^2+^ concentrations, respectively. **a:** Simulations for the same experimental setting as during optimization shows good agreement with the data. **b:** Model simulations successfully predict the results of new experiments. In addition to ON-switching with arabinose, the memory system was pushed into the OFF-state with increased Zn^2+^ concentrations. Shown here are the experimental and simulated densities for cells that were switched ON (Ara), OFF (Zn^2+^), and ON again. Densities for ON-OFF and ON-OFF-ON-OFF experiments can be found in Appendix Figure S5. The fluorescence intensities are shown on a logarithmic scale.

While the data were generally well described, there are small differences between our prediction and the experimental data of the repeat experiment (Figure 5a). Mainly, mCherry fluorescence at low concentrations shows less experimental variance than our simulation. Also, the drift to the OFF-state is slightly slower in the repeat experiment than in the previous one. Both effects are most likely due to small differences in the experimental procedures between the two datasets.

Predicting the experiments that include fast OFF-switching via an increased Zn^2+^ concentration via model simulations required adjusting the parameter *ũ*_2_, which determines how much an elevated Zn^2+^ concentration increases the ZnF4-DNA binding constant. Overall, the model reproduces the multiple ON-OFF-switches, but struggles to decrease the CcrM concentration and thus the EGFP fluorescence as rapidly as observed in the experiments (Figure 5b). This can be seen on day three, where the experimental population quickly reaches very low EGFP intensities, corresponding to the OFF-state, after the addition of Zn^2+^. The simulated population, on the other hand, catches up to these low CcrM concentrations only on day four. This discrepancy is not surprising, since our model calibration had no information on CcrM degradation rates in the steady state, which determine how quickly the CcrM concentration decreases after an abrupt stop of CcrM expression.

### 2.6 Optimization Reveals Two Possible Mechanisms for OFF-Drift

One question we wanted to explore with our model was when and how OFF-switches occur in individual cells. From our previous work and biological reasoning, we hypothesized that the faster cells divide, the faster they lose methylation, and therefore they should tend to switch back to the OFF-state sooner. To investigate this on a single-cell level, we colored the simulated EGFP trajectories by the average time between divisions (*T*_*div*_) for each cell (Figure 6a, f). This average differs between cells, because we generated the inter-division time samples for an individual cell and its daughter cell in a correlated manner to replicate inheritance effects.

**FIGURE 6.**
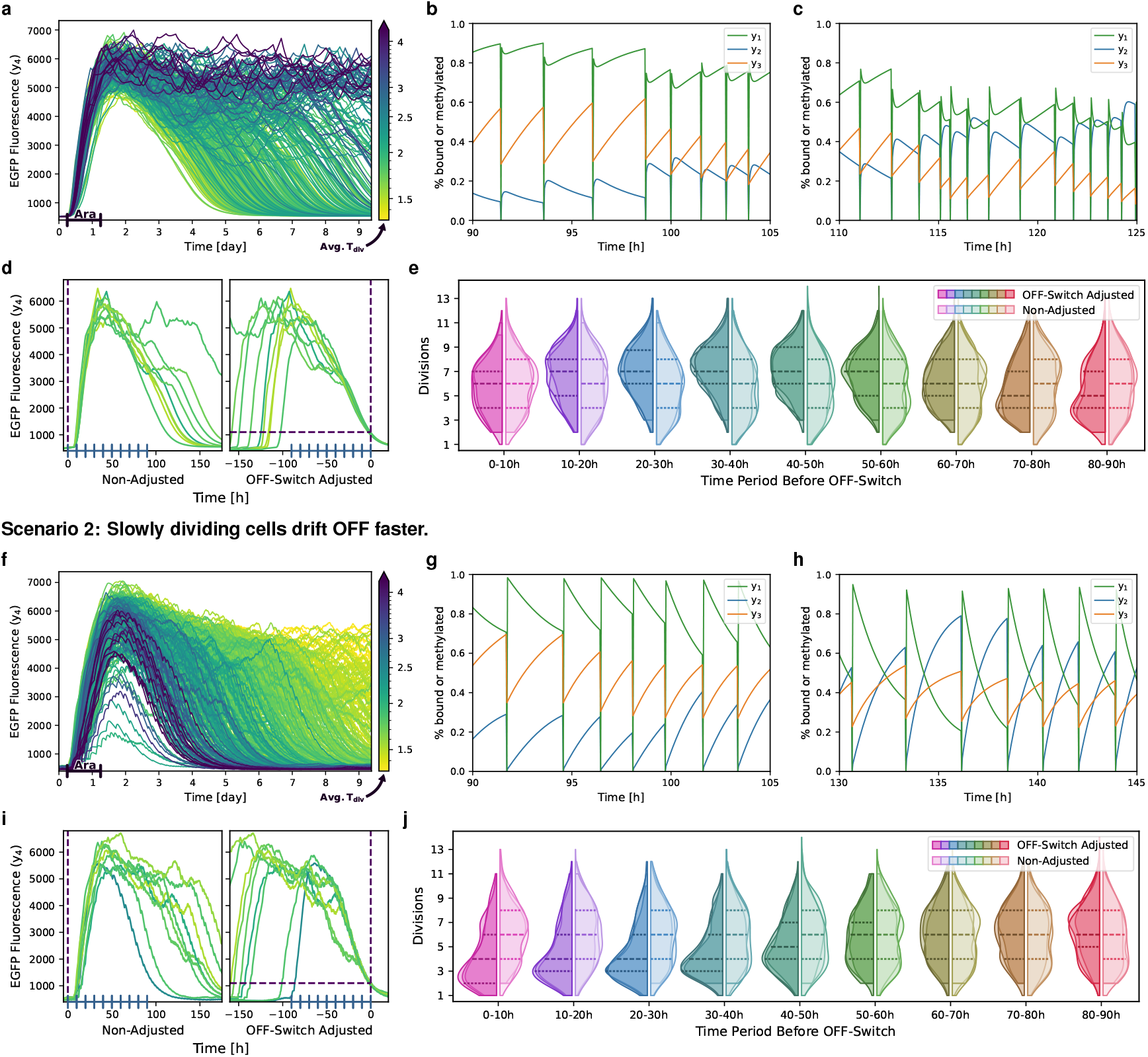
Two competing model scenarios. Plausible parameter values cluster into two groups that differ in how cell cycle length affects the ON-state stability of the memory system. **a–e:** In Scenario 1, the ON-state of quickly dividing cells is less stable. **f–j:** In Scenario 2, the ON-state of slowly dividing cells is less stable. Both scenarios were subject to the same analysis: **a+f:** Simulated single-cell trajectories of a population of cells upon induction with arabinose. Average cell cycle lengths are color coded. **b+g:** Exemplary state trajectories of a single cell in the ON-state, and **c+h:** exemplary state trajectories of a single cell close to switching to the OFF-state. The states *y*_1_ and *y*_2_ correspond to the degree of plasmids with bound CcrM or ZnF4, respectively, and *y*_3_ is the degree of methylation in one cell. **d+i:** To analyze the timing of the decision of an individual cell to switch to the OFF-state, we compared the number of cell divisions in the population in time intervals of 10 hours at different time points before the switch timepoint (OFF-switch adjusted, left) with the numbers for time intervals that were not adjusted for individual switching times (right). **e+j:** Result of this analysis. Shown is a comparison of observed distributions of average cell divisions between non-adjusted and OFF-switch adjusted time intervals of equal lengths of 10 hours. The distributions of three simulations with different sets of randomly sampled values are superimposed, with the dashed lines indicating the quartiles of the combined distributions.

During this analysis, we discovered two sets of parameters that showed opposite behavior in terms of the correlation between division rates and OFF-switching rates. We classified these two sets as those that behave according to our initial hypothesis (Scenario/Group 1, Figure 6a-e) and those that behave the opposite way, with the slowly dividing cells turning off faster (Scenario/Group 2, Figure 6f-j). Below we compare the two groups using a representative set of parameters from each group. Simulated trajectories, as in Figure 6a+f, for five parameter sets from each group can be found in Appendix Figure S2. In Figures 4 and 5 the group 2 parameter set was used; violin plots of the distributions for the group 1 parameter set can be found in Appendix Figure S4.

#### Stochasticity in Simulated Trajectories

In addition to their opposite division/switching correlation, the two groups also show other differences. The EGFP fluorescence intensities of group 1 cells (Figure 6a) were simulated as relatively smooth curves over time, interrupted by larger jumps. This indicates that the CcrM levels, and therefore EGFP intensities, do not vary much within a cell cycle, and the larger jumps are mostly due to changing plasmid numbers. The simulated trajectories of group 2, on the other hand, fluctuate more on a shorter time scale, giving the curves a wavering appearance (Figure 6f). These changes are on the time scale of cell divisions, so the simulated CcrM levels vary much more within each cell cycle than in group 1.

The cells in group 1 that remain in the ON-state for the ten days of the observation period remain on average at a constant fluorescence intensity, whereas in group 2 the intensity decreases slowly over time.

Furthermore, in group 1, all cells show a similar response to induction with arabinose and only begin to return to the OFF-state after four days. In group 2, there are a few cells that are only weakly responsive to the induction, and even cells that reach the ON-state can return to the OFF-state more quickly than those in group 1.

#### Close-Up Trajectories Reveal Interactions at DNA Level

The close-up trajectories for a sample cell from each group highlight the behavior over the course of a cell cycle that was already visible in the EGFP time courses (Figure 6b+c, g+h). The trajectories differ between parameter sets of the same group at a very detailed level, but the overall effects are similar within a group. Both bound CcrM (*y*_1_) and bound ZnF4 (*y*_2_) vary much more between cell divisions in group 2 than in group 1. This difference becomes more pronounced as the cells drift towards the OFF-state switch (Figure 6h). In both groups, methylation levels (*y*_3_) increase steadily during a cell cycle, as DNA replication during cell division is the only process that decreases methylation levels.

After a cell division, in which CcrM and ZnF4 are removed from the DNA by the DNA replication machinery, they compete for access to the DNA. There is a fast and a slow time scale in group 1 cells. On the short time scale, CcrM binds faster than ZnF4, followed by the delayed ZnF4 binding, which decreases the amount of bound CcrM by competition. On the slower time scale until the next division, bound CcrM increases the DNA methylation levels and therefore bound ZnF4 decreases, indicating a dynamic equilibrium between CcrM and ZnF4 binding to the DNA. In the ON-state, the overshoot of bound CcrM directly after a cell division is less than the level reached before the cell division, following the increased methylation levels (Figure 6b). During the drift towards the OFF-state switch, this initial overshoot of bound CcrM becomes larger, exceeding the levels of bound CcrM before the cell division (Figure 6c). This is the case because of increased competition by ZnF4, which can bind more strongly to the DNA with decreasing methylation levels.

In group 2, we do not observe such different time scales as in group 1. Here, ZnF4 binding is slower than CcrM, resulting in higher levels of bound CcrM directly after a division, which decrease slowly until the next division, while bound ZnF4 increases, indicating a competition of both proteins at unmethylated DNA sites. The increase in bound ZnF4 slows near the next division due to increasing DNA methylation levels, which block ZnF4 binding. These data also indicate that DNA methylation levels increase only slowly during the cell cycle. The decrease in bound CcrM, after the initially high levels following a cell division, becomes more rapid as the cell drifts toward the OFF-state (Figure 6h). This is again due to lower methylation levels leading to an increase in ZnF4 binding to the DNA and therefore more competition for a binding site, but also decreased levels of free CcrM competing with ZnF4.

#### Group Differences of Parameter Values

We identified the ZnF4 association (*a*_2_) and dissociation (*d*_2_) rates as the largest discriminator between the two groups at the level of parameter values (Table 2 and Appendix Table S2). In group 1, both rates are faster, i. e. ZnF4 binds to the DNA faster, but not as strong as in group 2. Especially in comparison to the CcrM dissociation rate *d*_1_, ZnF4 dissociates from the DNA much slower in group 2, indicating that in this setting a ZnF4 molecule once bound to the DNA can occupy the site for a long time and protect it from access of CcrM. Other parameters such as the CcrM production (*k*_4_) and degradation (*d*_4_) rates as well as the methylation rate (*k*_3_) are very similar across many optimizations and between the two groups. It is reasonable that they are well identifiable since *y*_3_ and *y*_4_ are the states for which we have direct experimental data. Parameters like CcrM association (*a*_1_) and dissociation (*d*_1_) rates vary more between different optimizations, but not discernibly between the two groups. The objective function values of the two groups are similar, but show a slightly better fit to the data for group 2.

#### Population Analysis of Switching Dynamics

We also wanted to identify the time period how long a cell divides faster or slower before switching into the OFF-state. This can give us an idea of when cells reach a point of no return, where they can no longer remain in the ON-state. We therefore analyzed how the distribution of division counts in a time interval of a fixed length changes over time. Since we are interested in the time period before a cell switches to the OFF-state, we set the reference point for these time intervals as the time when a cell falls below a threshold level of EGFP fluorescence. We will refer to this point as the switching time. We count the number of divisions for a series of tiled 10 h intervals, starting just before the switch occurs and going back in time to the beginning of the simulation. This process is illustrated in Figure 6d+i (right) for a small number of cells. As not all cells crossed the OFF-threshold during the simulation, and some crossed it very early, we only compared cells with a switching time between day four and day ten. As a reference for our comparison, we also counted the number of divisions in 10 h time intervals starting from the beginning of the simulation for the same cells. For these reference distributions, the time axis was not adjusted to the individual switching time (Figure 6d+i, left).

The resulting distributions of the numbers of divisions in 10 h time intervals before the OFF-switch and in the original order as reference are shown as violin plots for both groups (Figure 6e+j). There is an immediate difference between the two groups for the interval just before the OFF-switch (0–10 h). In group 1, the number of divisions follows the reference distribution, while in group 2, there are significantly fewer divisions. Going back in time, in group 1 the deviation from the reference distributions starts 10 h before the OFF-switch, with significantly more divisions, and is strongest between 30–50 h before the OFF-switch. At 60 h before the switch, the number of divisions returns to the reference distribution. In case of group 2, the reduced number of divisions continues until 60 h before the switch, with a maximum difference between 20– 40 h. In conclusion, in group 1, accelerated cell divisions start 50 h before the switch to the OFF-state, and the final irreversible push to the OFF-state occurs at least 10 h before the threshold is crossed. In group 2, reduced cell divisions also start about 50 h before the OFF switch, but they can also occur very close to the threshold. This difference between the groups of when the OFF-switch is certain can also be seen in the trajectories that are aligned to their switching time (Figure 6d+i, right). In group 1, all trajectories are lined up for the last 20 h, starting from EGFP fluorescence intensities of roughly 3000, while in group 2 they are aligned for only the last few hours before the OFF-threshold, starting from EGFP fluorescence intensities of less than 2000.

From this analysis, we cannot determine how many faster or slower divisions are needed, or whether they need to occur consecutively, to push a cell irreversibly into the OFF-state. However, for both groups we can infer that the effects of several shorter or longer divisions must accumulate to successfully destabilize the ON-state. This assumption is based on the fact that in both groups for a period of 20–30 h the number of divisions is significantly increased or reduced, as over 75% of the distributions are above or below the median for groups 1 and 2, respectively.

#### Which Scenario is Correct?

The two model scenarios and respective sets of parameters presented in the previous section can both explain the experimental data equally well. We defined them in terms of their OFF-switching behavior, namely whether OFF-switching is more likely for quickly or slowly dividing cells. However, our simulations of states at the DNA level, as well as the parameter analysis, allowed us to classify them according to their ZnF4 binding dynamics.

In group 1, the main effect of cell division is, as we previously hypothesized, that it leads to the reduction of DNA methylation and therefore weakens the ON-state. If the cells divide too fast, CcrM cannot keep up the pace to remethylate the DNA, allowing ZnF4 to bind more sites and inhibit further CcrM transcription in a self-enforcing process (Figure 7a, top).

**FIGURE 7.**
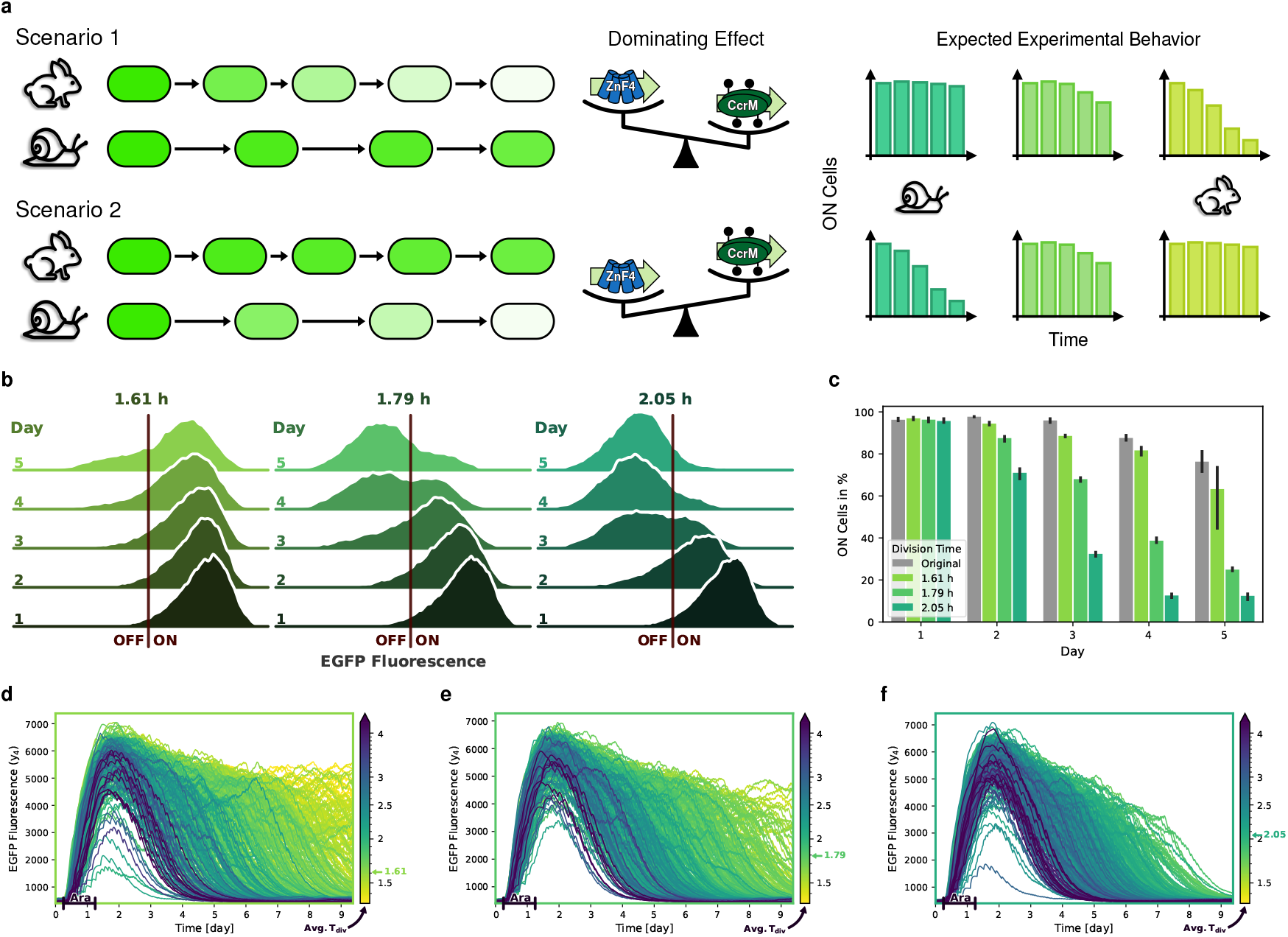
New experiment clearly favors scenario 2. **a:** Illustration of the two competing scenarios describing the effect of cell division rate on the stability of the ON-state. In scenario 1, the ON-state is expected to be less stable in quickly dividing cells (rabbits) compared to cells which divide more slowly (snails). Dilution of DNA methylation and CcrM binding is the dominating effect of cell division. Hence, the ON-state is expected to be more stable in cell populations that divide more slowly. In scenario 2, this is reversed, and the ON-state is predicted to be more stable in quickly dividing cells. Removal of ZnF4 is the dominating effect of cell division, and the ON-state is hence expected to be more stable in slowly dividing cells. **b-c:** Experimental results. **b**: EGFP fluorescence of cells in growth conditions with three different peptone concentrations: 10 g/L: 1.61 h generation time, 2.5 g/L: 1.79 h, 0 g/L: 2.05 h. Exponential fits to growth curves to determine the growth rate can be found in Appendix Figure S9. **c:** Percentages of cells in the ON-state in the three different growth conditions, as well as of the original experiments, over time. Cells with EGFP fluorescence intensities over the threshold, indicated in b), are considered to be in the ON-state. **d-f:** Single-cell trajectory simulations corresponding to three experimental conditions: **d:** Original division rates (average cell cycle length 1.61 h), **e:** more slowly dividing cells (average cell cycle length 1.79 h) and **f:** very slowly dividing cells (average cell cycle length 2.05 h).

In group 2, on the other hand, ZnF4 binds so strongly that it protects the DNA from methylation. Its dissociation from the DNA during DNA replication is an important factor needed to maintain DNA methylation and, therefore, DNA replication helps to prevent the switch to the OFF-state. Because CcrM can bind more quickly after DNA replication, a replication cycle allows DNA to be methylated again and more CcrM to be transcribed before it is replaced by ZnF4. Consequently, when cells divide slowly, more ZnF4 binds over time, protecting more and more target sites against methylation. The unmethylated DNA state supports ZnF4 binding in a self-enforcing process finally promoting the switch to the OFF-state (Figure 7a, bottom).

If scenario 1 is correct, we would expect more cells to remain in the ON-state if they divide more slowly and fewer if they divide more rapidly, and vice versa for scenario 2 (Figure 7a, right). In the following section, we investigate this question experimentally.

### 2.7 New Experiments Surprisingly Favor Scenario 2

To reduce the cell growth rate, ideally without otherwise affecting our system, we cultivated bacteria with a reduced peptone concentration (10 g/L vs. 2.5 g/L and 0 g/L). Lowering or raising the cultivation temperature was not appropriate as an interrogation to modulate cell growth, because this would also change the binding strengths of ZnF4 and CcrM to the DNA, as shown in earlier experiments (Maier et al. 2017). Also, a reduced glucose concentration in the medium could not be used, because this would weaken the inhibition of trigger CcrM expression. Restriction analysis confirmed that global DNA methylation levels were not altered by the reduced peptone concentrations, suggesting that any changes in the switching dynamics are indeed due to the reduced growth rate (See Appendix Figure S8).

We determined generation times of 1.79 h and 2.05 h for the cultures with 2.5 g/L and 0 g/L peptone, respectively, compared to the generation time of 1.61 h for the culture with the original 10 g/L peptone medium (Appendix Figure S9).

Cultivation of cells under these three conditions over five days resulted in a surprisingly clear difference in ON-state stability (Figure 7b). While a large percentage of cells at the original growth rate stay in the ON-state over five days, cells at lower growth rates switched to the OFF-state much sooner. The percentage of cells in the ON-state, as indicated by fluorescence intensities above the threshold (red line), is quantified in Figure 7c, including the ONpercentages of the original experiments (gray). The population with a generation time of 1.61 h starts to drift into the OFF-state about one day earlier than in our previous experiments. However, the relative changes to the populations with reduced growth rates can still be compared to our model simulations. While over 60% of the cells with the fastest growth rate remain in the ON-state after five days, less than 25% of the cells with a generation time of 1.79 h remain ON and fewer than 15% for cells with 2.05 h generation time. One day after the arabinose induction was removed (Day 2), only 70% of the cells at the slowest division rate are still in the ON-state, and only half of these on day three. This indicates that at these low division rates, cells start to switch OFF directly after termination of the arabinose induction. Cells with the intermediate generation time (1.79 h) reach the same ratio of OFF-cells approximately one day later. The speed at which cells switch into the OFF-state therefore correlates with slower division rates. These observations clearly confirm the experimental behavior expected from model scenario 2, where more slowly dividing cells switch off faster.

Model simulations with inter-division times adapted to the experimental generation times show the same qualitative behavior (Figure 7d-f). At the slowest division rate, the simulated EGFP fluorescence decreases sharply after the induction ends and all cells are in the OFF-state at the end of the simulation. The simulation, however, differs from the experimental data in the speed at which cells switch off. Though, such a discrepancy was to be expected, as it is plausible to assume that all parameters will slightly change upon fundamental alterations of the growth conditions.

In summary, our model has led to two competing scenarios regarding the role of cell division in the stability of the ON-state. Surprisingly, subsequent experiments examining this stability in cell populations with different average cell cycle lengths revealed a positive correlation between cell division speed and ON-state stability, meaning that slowly dividing cells show a lower ON-state stability. Furthermore, our model suggests that ZnF4 binding is strong enough to prevents DNA methylation between cell divisions. Immediately after DNA replication, CcrM binds much faster to the DNA than ZnF4, thus providing a window of opportunity to maintain methylation levels after replication and thereby support the ON-state.

## 3 DISCUSSION

Epigenetic systems are attractive regulatory paradigms in synthetic biology and have received increasing attention in recent years. In this study, we have developed a population model for synthetic epigenetic memory systems (Figure 1). This was a challenging task for several reasons: (i) The system involves intertwined processes that occur on multiple time and size scales. First, biochemical reactions involve single molecules, such as the mutual inhibition of CcrM and ZnF4 binding or methylation events, and occur on a scale of minutes. Second, a cell divides every 2-3 hours and cell division affects the molecular state of the system as described. And third, the OFF drift of the population happens on a time scale of several days. (ii) Not all of these processes can be directly observed in experiments. We have direct experimental data for CcrM at the single-cell level and bulk DNA methylation data, both on the slow time scale. This means that we have to solve an inverse problem to infer parameters for the faster time and size scales that are able to reproduce the observed behavior of the memory system on the slow time scale.

The challenge is further complicated by the fact that the number of cells in the population is large.

Our aim was to describe the population behavior upon induction and its slow drift toward the OFF-state within nine days after the trigger is removed. Our hypothesis was that this slow drift of the population from ON to OFF is caused by the accumulation of stochastic events, in particular the stochasticity introduced by faster or slower cell divisions. Since the number of molecules involved in the process is large, we used a hybrid modeling approach, in which the biochemical processes in a single cell are described deterministically by chemical reaction kinetics, and cell division is described as a stochastic event that interferes with this reaction system. The population consists of a set of independently simulated cells (Figure 3) and can quantitatively mimic the single-cell dynamics of the CcrM expression, as well as the evolution of the bulk DNA methylation levels (Figure 4).

Parameter estimation was robust and reproducible, as many of the model parameters could be estimated with low variance despite different initial starting conditions and the stochastic nature of the model, which can be interpreted as suggesting that we have likely captured the main processes that characterize the behavior of the population. Our model was further validated by its ability to describe unseen experimental data generated in novel experimental workflows (Figure 5) and was then used to generate hypotheses about unobservable molecular interactions at different time scales.

The estimated parameters cluster into two groups that only differ slightly in terms of objective function values. However, the two groups behave opposite with respect to the influence of cell division duration on the stability of the ON-state after removal of the input trigger (Figure 6). In group 1, the ON-state of slowly dividing cells is more stable than that of rapidly dividing cells. This model was expected, as cell division leads to the reduction of DNA methylation. The opposite is true for group 2, where slow cell division is a driver for the switch from the ON to the OFF-state.

Experiments in which average cell growth rates were varied by cultivating cells under different nutritional conditions clearly support the behavior predicted in group 2, as the percentage of cells in the ON-state decreased significantly faster in populations with lower growth rates (Figure 7). Single-cell simulations and parameter analysis provided an explanation for this observation. According to the model, ZnF4 binding to the DNA is so strong that it prevents DNA methylation between cell divisions. During DNA replication, ZnF4 and CcrM are both dissociated from the DNA, but CcrM binds more rapidly immediately after the replication fork passage, allowing to maintain higher methylation levels over the time scale of many cell divisions. In addition, statistical analysis of the number of cell divisions in time intervals before the OFF-switch suggests that the decision to switch requires some accumulation of longer cell cycle lengths, which is already visible at the population level 40– 50 h before the switches occur in individual cells, and is most pronounced 20–30 h before the switches. Hence modeling combined with tailored experiments were able to document an unexpected and at first sight apparently illogical effect of stochasticity in cell division rates on the stability of the ON-state of the memory system investigated in our work.

By resolving snapshot data into individual state trajectories, our model allows hypotheses to be made about detailed interaction dynamics at the single-cell level. These hypotheses can drive further experiments to test these hypotheses, as demonstrated here. This has led to a more complete picture of the underlying processes at the DNA level that cannot be directly observed in experiments. According to this, the data can be explained by a tight binding of ZnF4 to the DNA, which efficiently prevents methylation between cell divisions. During DNA replication, both ZnF4 and CcrM are dissociated from the DNA, and immediately after replication, CcrM rebinds more rapidly than ZnF4, but not as strong in the long term. Thus, rapid division promotes CcrM binding and subsequent methylation over the longer time scale of days. This picture was a result of the modeling process and has revised our previous belief about the role of cell division in ON-state stability. In particular, one of our very early model assumptions was that CcrM binding was the rate-limiting step and methylation occurred immediately after binding. This assumption was revised during the model fitting process (Figure 2), where we decided to decouple methylation from CcrM binding to the DNA, which significantly improved our model fits.

Our model also sheds light on the mechanisms behind the drift of the population back to the OFF-state. According to our picture, the mechanisms of the switch from the OFF- to the ON state upon induction with arabinose and the drift back to the OFF-state are fundamentally different. CcrM from the trigger plasmid shifts the binding equilibrium of the competing ZnF4 and CcrM strongly towards CcrM, thus turning on the positive feedback loop. This happens on a fast time scale and is not much affected by stochastic cell division. In contrast, the slow drift of the population towards the OFF-state is driven by an accumulation of stochastic effects, leading to self-enforcing processes that cause an increasing probability over time for cells to switch. This process cannot be captured by deterministic simulations. After the triggering phase, the ON-state is a transient metastable state with a long average residence time. These residence times show a high variability across cells, and the observed slow drift and transient bimodal behavior at the population level are collective effects of these individual residence times. Conceptually, this is similar to noise-driven unlimited population growth as described for example in Meerson and Sasorov (2008), where the deterministic system described by reaction rate equations converges to a stable equilibrium on a fast time scale, which becomes metastable in stochastic simulations, and the population leaves the metastable state and grows unlimited on a longer time scale.

The challenges that we faced for the modeling of this system are similar to the simulation challenges of many other biologically relevant systems. These include for example, the stimulation of specific signaling pathways in cell populations under control and perturbation conditions and induced phenotypic responses (Santos et al. 2007, Zhang et al. 2014) or metastable switches in synthetic biology, pioneered by the development of the “toggle switch” and “repressilator” systems (Elowitz and Leibler 2000, Gardner et al. 2000). The hybrid modeling approach we have developed thus may provide a valid template for many other settings, allowing the investigation of phenomena such as metastability and slow drifts that cannot be explained by purely deterministic approaches.

Our modeling approach is, of course, based on simplifying assumptions. Possible extensions of the model could address the most drastic simplifications and, depending on the goals and questions, include for example the effect of changes of cell volume during the cell cycle on molecule concentrations and the partitioning of contents to daughter cells. In addition, in a real cultivation experiment we would expect the proportion of rapidly dividing cells to increase over time, which is not accounted for in our model because we follow only one daughter cell to keep cell numbers constant. We also do not account for a putative selective advantage or disadvantages, for example of other methylation events catalyzed by the exogenous DNA methyltransferase CcrM on the *E. coli* genome, which for example can affect bacterial gene regulation (García-Pastor et al. 2019).

In conclusion, we believe that our combined approach of hybrid modeling and experimentation can serve as a template for a general description of epigenetic processes and, more generally, metastable systems in synthetic biology as well as in nature.

## 4 METHODS

### 4.1 Probability Distributions and Sam-pling

Our model includes different stochastic elements to account for the observed heterogeneity in the experimental data.

#### Copy number fluctuations

The constant *Z*_*P*_ captures the stochasticity in memory plasmid copy numbers per cell and its influence on the abundance of ZnF4 (*Z*_*Z*_). For every cell and after every division it is sampled randomly from a Beta distribution. The parameters of the distribution are based on experimental observations resulting in Beta(*α* = 4, *β* = 4), shifted and scaled to range from 90–230, with a mean of 160.

#### Varying induction and repression strength

The constant *Z*_*T*_ represent the combined stochasticity resulting from copy number fluctuations of the trigger plasmid as well as variations in the strength of the induction or repression of the arabinose operon. This value is sampled only once per cell, as the trigger subsystem is only important for a short period of time. *Z*_*T*_ follows a lognormal distribution with *µ* = 0 and *σ* = 0.3.

#### Measurement noise

The distributed parameters were not sufficient to describe the variance observed in the measurements, especially at low fluorescence intensities, corresponding to no or very little expression of EGFP or mCherry. We therefore added measurement noise to the simulation data before comparison to the experimental data. This was only done for population data, not for individual time courses. The noise was sampled from a normal distribution with *µ* = 0 and *σ* = 0.6 for EGFP and *σ* = 0.25 for mCherry and then scaled with value of the simulated intensity giving: *y*_{4,5}, noise_ = *y*_{4,5}_ + *Z*_*n*_ *· y*_{4,5}_. See Appendix Section 2 for more detail.

#### Stochastic division events

A stochastic process captures the stochasticity of cell division timing. The time between each event, *T*_*div*_, is randomly distributed and follows an exponentially-modified normal distribution. The probability density

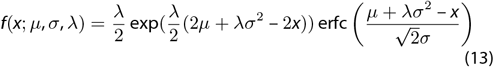

is the convolution of the densities of a normal and an exponential distribution. The parameters were chosen, so that the mean resembles the generation times obtained from growth curves. These are *σ* = 0.17, *λ* = ^1^/_0.935_ and *µ*_1.61_ = 0.675, *µ*_1.79_ = 0.855, *µ*_2.05_ = 1.115 for cells with average generation times of 1.61, 1.79, and 2.05 respectively.

As more quickly dividing cells are more likely to also divide quickly in future and vice versa for more slowly dividing cells, we replicated this inheritance effect by generating correlated samples for the inter-division times. This was done via Markov chain Monte Carlo (MCMC) sampling, where each sample chain produces the inter-division times for one cell, which are therefore correlated over time.

### 4.2 Simulation Framework

All programming was done in Python 3.10 and 3.12, using Numpy (Harris et al. 2020), Scipy for statistics tools (Virtanen et al. 2020), Pandas for dataframe handling (McKinney et al. 2010), and Matplotlib (Hunter 2007) and Seaborn (Waskom 2021) for creating graphics. The model was defined and simulated with Tellurium (Choi et al. 2018, Medley et al. 2018). Model files are available in the human-readable format antimony used by Tellurium, as well as in SBML format. Computationally intensive parameter optimizations were run on the high-performance cluster bwUniCluster (2.0).

### 4.3 Optimization

**ZnF4 Penalty** We calculate the distance for all cells of the ratio of bound ZnF4 (*y*_2_) to the threshold

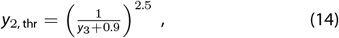

where *y*_3_ is the methylation level. The penalty then is the sum of all distances with values above the threshold (*Pen*_*ZnF*_). It is only calculated for the first two days, directly before and after induction, because we do not have data of this relationship from long-term experiments.

#### Objective Function Weights

The weights in the objective function were chosen such that all terms are of similar magnitude, but with *Kol*_*fcm*_ having the most significant impact:

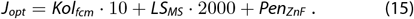

#### Optimization Algorithm and Procedure

The global optimization algorithm differential evolution from Scipy (Virtanen et al. 2020), which is based on Storn and Price (1997), was used to calibrate the model parameters to the experimental data. It is a stochastic algorithm and does not require a gradient, which would be difficult to calculate accurately due to stochastic nature of our simulation. For the same reason, we also adjusted the tolerances for the objective function values, which define the stopping criterion of the algorithm. Our aim was to allow a variation in the objective function in the order of the variation observed, when simulating the same parameter set repeatedly.

As the optimization problem is computationally expensive and can have multiple similarly good solution, due to the stochastic components, we calibrated the model step-wise. Initially, we used wide bounds on the parameters and optimized multiple times. From these results we constrained the bounds further, aiding in converging to a good solution more quickly, while taking care not to cut out another solution space.

### 4.4 Experiments

#### Analysis of ON-state stability at different growth conditions

To monitor the stability of the epigenetic memory system in the ON-state, cultivation of *E. coli* XL1-blue cells containing the components of the memory system was performed in LB medium with peptone concentrations of 0 %, 0.25 % and 1 % (w/v) in each case supplemented with 0.2 % glucose, 10 µm ZnSO4 and antibiotics at 28 °C as described in Ullrich et al. (2020), Klingel et al. (2022), Graf et al. (2023). To determine the doubling time of the cultures, the OD_600_ was measured over a period of 10 h for cultures cultivated in each of the three LB media in a separate experiment without induction. Induction was performed with the ON-state trigger medium (25 µg/mL kanamycin, 100 µg/mL ampicillin, 10 µm ZnSO4 and 0.02 % (w/v) arabinose) on the first day of the experimental series, and ON-state stability was monitored over 5 days by flow cytometry analysis as described in Ullrich et al. (2020), Graf et al. (2023).

## Data Availability

All model code, including the experimental data sets, sampled parameters and optimization results can found at doi: 10.15490/fairdomhub.1.model.871.1. The collection includes details of the modeling environment, a short description of all code files and which ones were used to create the figures.

## Additional Information

The Appendix includes in-depth information on modeling details, such as the mapping of fluorescence intensities to protein amounts, measurements noise, and the objective function. It also contains additional parameter sets and simulations, analogously to the main text, as well as the raw data on the objective function variability. It further includes the simulations without the different sources of heterogeneity in the model, and, lastly, the calculation of growth rates and data of methylation levels under different conditions.

## ACKNOWLEDGMENTS

Funded by DFG grants JE 252/35-1 (AJ) and RA 1840/2-1 (NR). Funded by Deutsche Forschungsgemeinschaft (DFG, German Research Foundation) under Germany’s Excellence Strategy EXC 2075 – 390740016. We acknowledge the support by the Stuttgart Center for Simulation Science (SimTech). The authors acknowledge support by the state of Baden-Württemberg through bwHPC.

## AUTHOR CONTRIBUTIONS

VK: conceptualization, formal analysis, investigation (modeling) methodology, software, visualization, writing – original draft, writing – review & editing; DG: Investigation (experiment), methodology (experiment), writing – review & editing; SW: Investigation (experiment), methodology (experiment), supervision, writing – review & editing; AJ: conceptualization, funding acquisition, investigation (experiment), methodology (experiment), supervision,visualization, writing – original draft, writing – review & editing; NR: conceptualization, funding acquisition, formal analysis, investigation (modeling), methodology (modeling), supervision, visualization, writing – original draft, writing – review & editing;

